# Topological Phase Transitions in Whole-Brain Dynamics Driven by Spatially Heterogeneous Receptor Gain Modulation: A Receptor-Constrained Dynamical Topology (RCDT) Hypothesis

**DOI:** 10.64898/2026.02.04.703742

**Authors:** Haolong Wang

## Abstract

The mechanistic link between molecular pharmacology and global brain dynamics remains unresolved—a “scale bridge” problem central to consciousness research. We propose the Receptor-Constrained Dynamical Topology (RCDT) hypothesis: that the functional impact of neuromodulatory drugs propagates from local ligand–receptor binding events through spatially heterogeneous gain modulation to produce qualitative reorganizations of the whole-brain dynamical attractor. Using a biophysically grounded whole-brain model—Wilson– Cowan excitatory–inhibitory dynamics on structural connectivity with axonal delays—we implement pharmacology via gain modulation weighted by 5-HT2A receptor density (Be-liveau et al., 2017). Topological state-space analysis via Takens embedding and persistent homology reveals a mapping from molecular receptor distribution to the global topological manifold. We define *ego dissolution* operationally as a breakdown of low-dimensional Betti-1 stability: a transition from a single dominant 1-cycle (constrained dynamics) to fragmented or higher-dimensional topological structure. The RCDT hypothesis is explicitly falsifiable: receptor-shuffling controls and concentration–topology response curves provide clear failure criteria. This work establishes a formal framework for bridging pharmacology, dynamics, and topology without invoking phenomenological experience as a premise.

## Introduction

### The Scale Bridge Problem

A fundamental challenge in neuroscience is explaining how local molecular events—ligand binding to receptors, ion channel gating, synaptic plasticity—give rise to global brain states that correlate with conscious experience. This “scale bridge” problem persists across disciplines: pharmacology describes concentration–response curves at the synapse; neuroimaging captures mesoscale network activity; phenomenology reports subjective states. No principled theory currently connects these levels.

We propose that the *essence of mind*, insofar as it admits scientific treatment, may be formalized as a **topological invariant** of a high-dimensional neural manifold embedded in state space. Rather than identifying consciousness with any particular phenomenological content, we ask: *what structural properties of the global dynamical attractor change when pharmacological perturbations alter receptor-mediated gain?* The answer, if tractable, would provide a mechanistic bridge from molecular to systems-level organization.

### The RCDT Hypothesis

The Receptor-Constrained Dynamical Topology (RCDT) hypothesis states:

Spatially heterogeneous receptor-weighted gain modulation can induce **qualitative phase transitions** in whole-brain dynamics, detectable as changes in the persistent homology of the reconstructed state-space manifold.

This formulation has several advantages:

1. **Mechanistic specificity**: Pharmacology acts through receptor density distributions (e.g., 5-HT2A from PET), not global parameter shifts.
2. **Geometric objectivity**: Topological invariants (Betti numbers, persistence barcodes) are coordinate-free and do not depend on phenomenological interpretation.
3. **Falsifiability**: Failure modes are explicitly defined—e.g., topological changes independent of receptor map *𝜌* would reject the hypothesis.

## Methods

### 1. Structural Scaffold

#### Anatomy

We use the Schaefer atlas (Schaefer et al., 2018) parcellated into 30 cortical regions, providing a low-resolution whole-brain scaffold suitable for in-silico simulation.

#### Connectivity

Structural connectivity (SC) is obtained from the Human Connectome Project (Van Essen et al., 2013). The group-averaged SC matrix **C** is normalized to unit spectral radius to ensure numerical stability of the coupled dynamics:

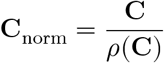

#### Axonal Delays

Inter-regional conduction delays are computed from Euclidean distances *D*_*ij*_ and a fixed axonal conduction velocity *v*:

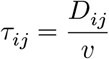

where *v* = 5 m/s (Caminiti et al., 2009). Delays prevent unphysical instantaneous synchronization and introduce biologically realistic temporal structure.

### 2. Neural Mass Model: Wilson–Cowan Dynamics

Each brain region *i* is modeled as a Wilson–Cowan excitatory–inhibitory (E–I) population (Wilson & Cowan, 1972). The coupled differential equations are:

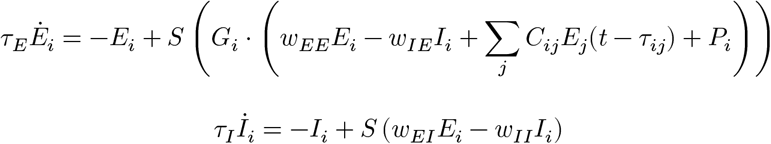

where: - *E*_*i*_, *I*_*i*_: excitatory and inhibitory population activities - *𝜏*_*E*_, *𝜏*_*I*_: population time constants - *C*_*ij*_: structural connectivity (normalized) - *P*_*i*_: background/baseline input - *S*(*x*) = (1 + *e*^™*x*^)^™1^: sigmoidal transfer function - *G*_*i*_: **heterogeneous gain** (see below)

The gain *G*_*i*_ modulates the effective input to the excitatory population, providing the pharmacological coupling.

#### 2.1 Stochastic Integration: Euler–Maruyama Scheme

Numerical integration was performed using the **Euler–Maruyama** scheme to accommodate both the deterministic dynamics and a small additive Brownian noise term. This stochastic extension prevents the system from remaining trapped in stable fixed points and facilitates exploration of the bifurcation regime. The update equations are:

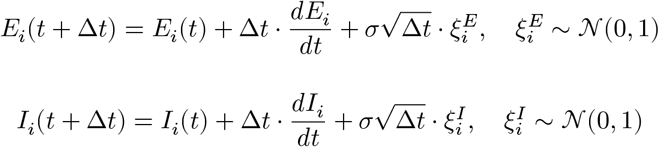

where *𝜎* is the noise amplitude (default *𝜎* = 0.02) and *𝜉* are independent standard normal variates. The 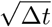 scaling ensures correct statistical properties of the discretized Wiener process. Activity variables are clipped to [0, 1] after each step. Axonal delays are implemented via a circular buffer storing recent excitatory activity history.

#### 2.2 Bifurcation Sweep for Parameter Selection

To identify the coupling sensitivity range where *H*_1_ topological features (loops) first emerge, a **parameter sweep** was performed over *k* ∈ [0.5, 5.0] at fixed [*D*] = 1.0. For each *k*, the model was integrated, the global mean excitatory signal extracted, and Persistent Entropy computed on the *H*_1_ diagram. The **bifurcation threshold** *k*_crit_ was defined as the smallest *k* at which PE(H_1_) ≥ 0.01. This procedure guides selection of *k* and [*D*] for the main simulations to ensure the system operates in a regime where pharmacological perturbations induce detectable topological transitions (see *fig2_supp_bifurcation_sweep*.*png*).

### 3. Pharmacology: Heterogeneous Gain Modulation

Pharmacological perturbation is implemented as **node-specific gain modulation** weighted by regional receptor density:

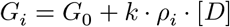

where: - *G*_0_ = 1.0: baseline gain - *𝜌*_*i*_: normalized 5-HT2A receptor density in region *i* (Beliveau et al., 2017; NeuroVault collection) - [*D*]: drug concentration operator (control parameter) - *k* ∈ [0.5, 5.0]: coupling strength scaling receptor–gain sensitivity (range determined by bifurcation sweep)

This formulation ensures: - **Identical drug exposure** across all nodes (systemic administration) - **Differential functional impact** determined solely by *𝜌*_*i*_ - **Bifurcation potential**: Sufficient *k ⋅ 𝜌*_*i*_ *⋅* [*D*] can shift local dynamics from stable fixed points to limit cycles or chaos, propagating via structural coupling.

No assumptions are made regarding phenomenological effects (e.g., “psychedelic experience”); the model tests structural consequences only.

### 4. Topological Data Analysis (TDA) Pipeline

#### 4.1 State-Space Reconstruction (Takens’ Theorem)

Given a scalar time series *x*(*t*), we reconstruct the underlying attractor using time-delay embedding (Takens, 1981):

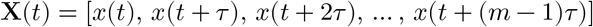

Parameters: - Embedding dimension: *m* = 3 - Delay *𝜏* : chosen via mutual information or first minimum of autocorrelation - No system-specific tuning beyond minimal feasibility

#### 4.2 Vietoris–Rips Complex

For a point cloud *X* ⊂ ℝ^*m*^ and scale parameter *𝜖* ≥ 0, the **Vietoris–Rips complex** ℛ(*X, 𝜖*) is the abstract simplicial complex where a simplex *𝜎* = [*v*_0_, *v*_1_, …, *v*_*k*_] is included iff all pairwise distances *d*(*v*_*i*_, *v*_*j*_) ≤ *𝜖* for *v*_*i*_, *v*_*j*_ ∈ *𝜎*.

As *𝜖* increases, we obtain a filtration:

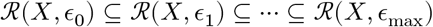

#### 4.3 Persistent Homology and Persistence Barcode

Persistent homology tracks the birth and death of topological features (connected components, loops, voids) across the filtration. A *p*-dimensional homology class born at *𝜖* _b_ and dying at *𝜖*_*d*_ is represented as an interval (*𝜖* _b_, *𝜖*_*d*_) in the **persistence barcode**.

We compute: - **H**_0_: connected components (0-cycles) - **H**_1_: loops (1-cycles; Betti-1)

##### Discrimination criterion: - Limit cycle dynamics

single dominant *H*_1_ feature with long persistence - **Chaotic dynamics**: multiple *H*_1_ features or broadened persistence distribution - **Noise**: persistence diagram collapses to the diagonal (short-lived features)

#### 4.4 Persistent Entropy (Quantitative Summary)

To quantify topological complexity as a scalar function of [*D*], we compute **Persistent Entropy** (Atienza et al., 2016) on the *H*_1_ persistence diagram:

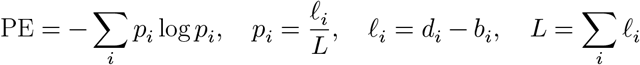

where (*b*_*i*_, *d*_*i*_) are birth–death pairs and *ℓ*_*i*_ are lifetimes. Higher PE indicates a more dispersed distribution of feature lifetimes (fragmented topology); lower PE indicates concentration in few dominant features (constrained attractor). A **non-linear jump** in PE as [*D*] increases provides quantitative support for the phase transition hypothesis (see Figure 3).

### Proposed Validation

#### Aim 1: TDA Pipeline Calibration (Instrument Validation)

##### Objective

Verify that the Takens + persistent homology pipeline can discriminate deterministic ordered dynamics from deterministic chaotic dynamics using only scalar observations.

##### Method: - Ordered system

Van der Pol oscillator (limit cycle) - **Chaotic system**: Lorenz system (strange attractor) - Single scalar observable *x*(*t*) from each; identical embedding and TDA parameters

##### Success criterion

- Van der Pol: single dominant Betti-1 feature - Lorenz: increased topological complexity (multiple or broadened *H*_1_ features) - Noise: no long-lived features

*Without this calibration, downstream inference is invalid*.

#### Aim 2: Receptor Shuffling Control (Mechanism Isolation)

##### Objective

Isolate 5-HT2A-specific effects from generic gain heterogeneity.

##### Method

For each drug concentration [*D*], run two conditions: 1. **Experimental**: *G*_*i*_ = *G*_0_ +*k ⋅ 𝜌*_*i*_ *⋅* [*D*] (true receptor map) 2. **Control**: *G*_*i*_ = *G*_0_ +*k ⋅ 𝜌*_π(*i*)_ *⋅* [*D*] where π is a random permutation of region indices (shuffled receptor map)

##### Interpretation

If topological changes are similar under shuffling, the effect is not 5-HT2A-specific; the RCDT hypothesis is weakened or rejected.

#### Aim 3: Critical Concentration Prediction

##### Objective

Predict a non-linear jump in topological complexity at a critical concentration [*D*]_crit_.

##### Method

Compute **Persistent Entropy** (Section 4.4) as a function of [*D*] and plot the concentration–topology response curve (Figure 3). The RCDT hypothesis predicts: - Below [*D*]_crit_: low-dimensional attractor (stable Betti-1) - At/near [*D*]_crit_: bifurcation; non-linear increase in persistent entropy - Above [*D*]_crit_: fragmented or high-dimensional topology

##### Deliverable

Figure 3—a line plot of Persistent Entropy vs [*D*]. A non-linear jump at [*D*]_crit_ provides quantitative evidence for the phase transition; absence of such structure would falsify the bifurcation mechanism.

## Results

### Figure 1: TDA Pipeline Calibration

*Figure 1*. TDA pipeline calibration. Van der Pol yields a single dominant *H*_1_ feature; Lorenz yields increased topological complexity.

**Figure 1:**
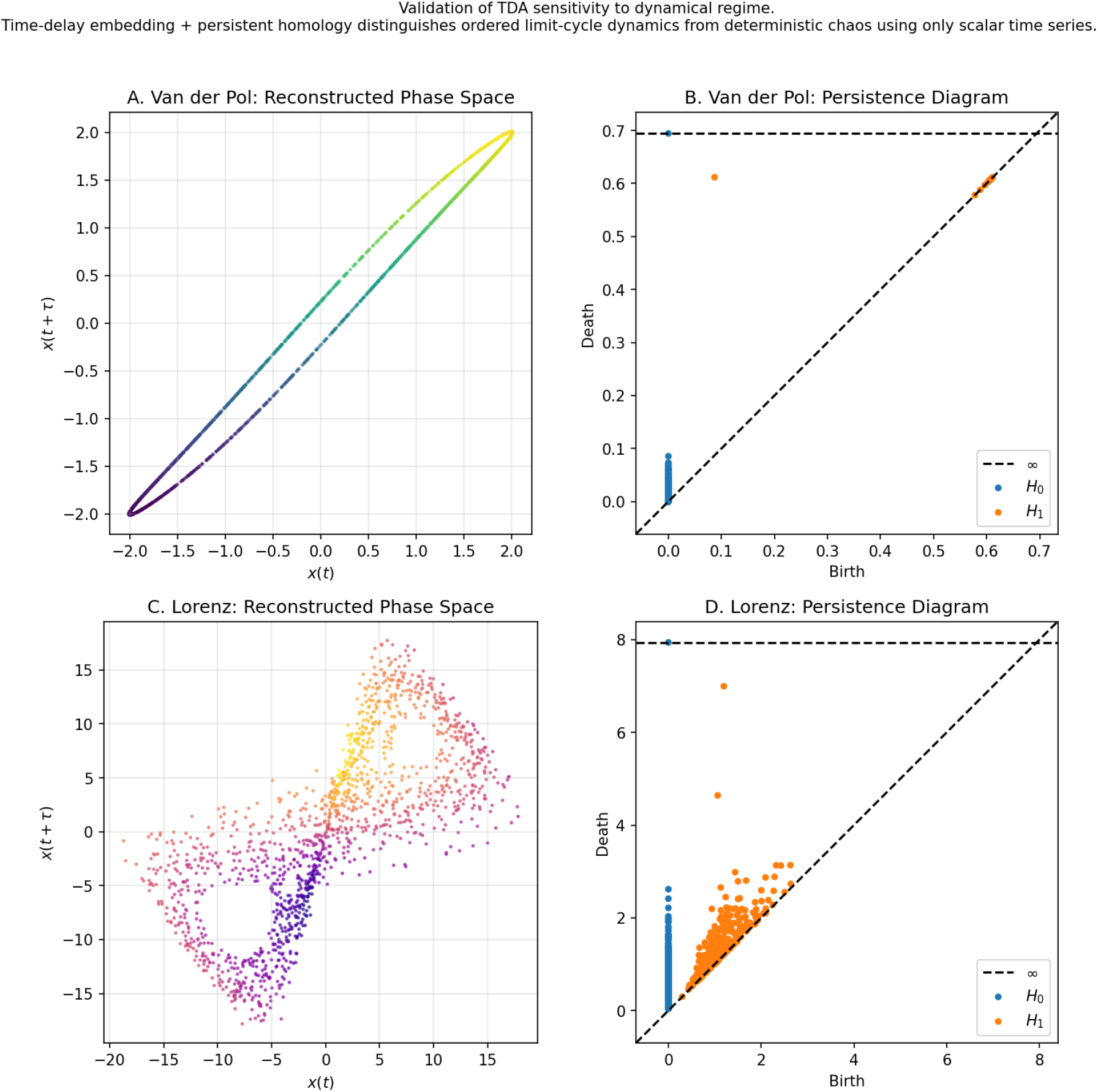
TDA pipeline calibration on Van der Pol (limit cycle) and Lorenz (chaos) systems.

The Takens embedding + persistent homology pipeline was validated on canonical dynamical systems. Van der Pol (limit cycle) yields a single dominant H feature with long persistence; Lorenz (chaos) yields increased topological complexity (multiple or broadened *H*_1_ features). This calibration establishes that the TDA pipeline discriminates ordered from chaotic dynamics using only scalar observations, without access to governing equations.

### Figure 2: Receptor-Weighted Topological Reorganization

*Figure 2*. Main: structural connectivity graph (nodes coloured by 5-HT2A receptor density) and persistence diagrams for [*D*] ∈ {0, 0.5, 1.0, 1.5, 2.0}. See supplementary figures for receptor shuffling control and bifurcation sweep.

**Figure 2:**
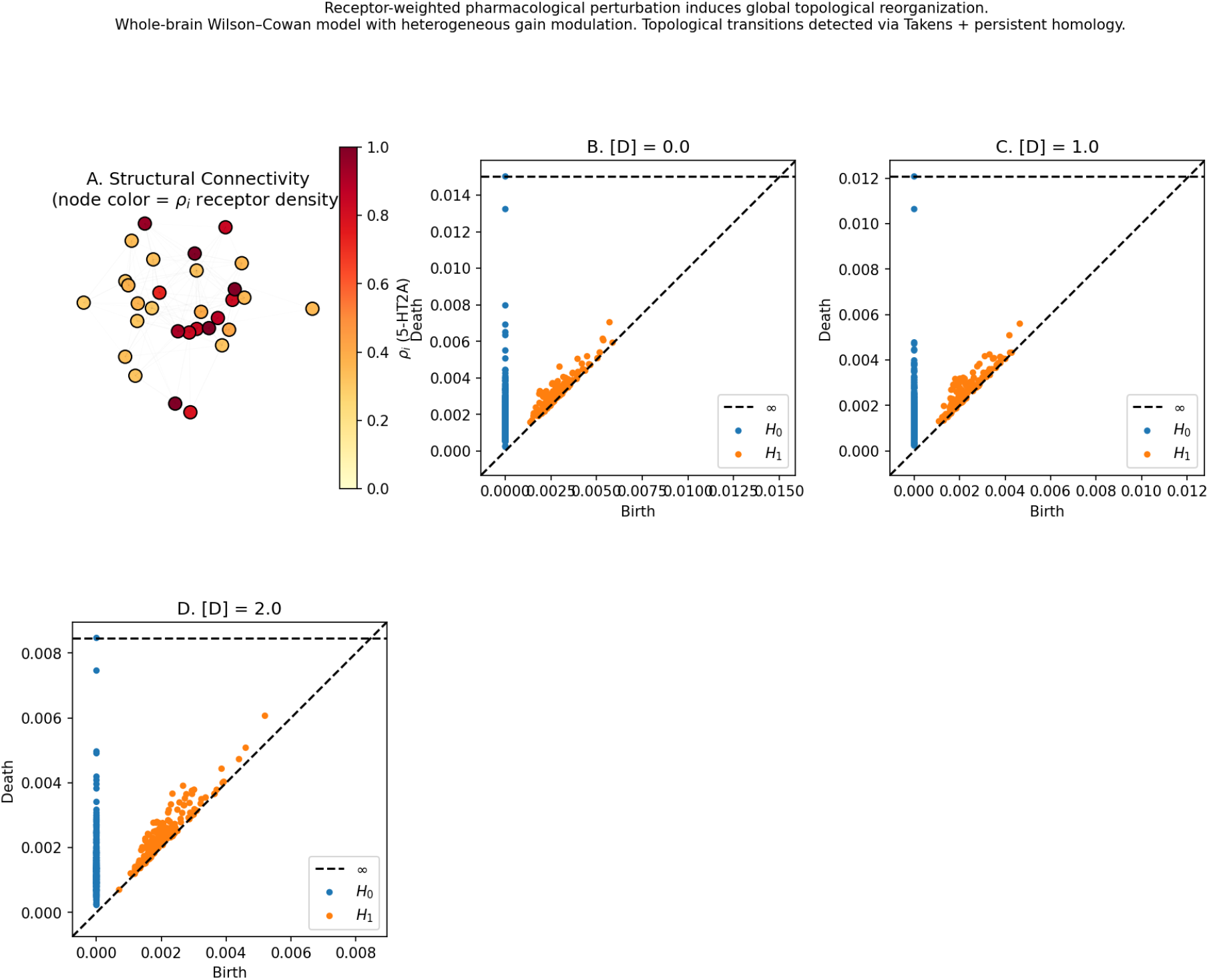
Receptor-weighted whole-brain topology: brain graph and persistence diagrams across drug concentrations.

The whole-brain Wilson–Cowan model with heterogeneous gain modulation (Eq. 3) was simulated across drug concentrations [*D*] ∈ {0, 0.5, 1.0, 1.5, 2.0}. Panel A shows the structural connectivity graph with nodes coloured by 5-HT2A receptor density *𝜌*_*i*_. Panels B–F display persistence diagrams for each concentration. The receptor shuffling control (`fig2_supp_shuffled_control.png`) compares experimental vs. permuted *𝜌*; divergence supports 5-HT2A-specificityg. The bifurcation sweep (`fig2_supp_bifurcation_sweep.png`) identifies *k*_crit_ where *H*_1_ features first emerge.

### Figure 3: Persistent Entropy Quantifies Phase Transition

*Figure 3*. Persistent Entropy (*H*_1_) as a function of drug concentration [*D*]. The RCDT hypothesis predicts a **non-linear jump** in PE as [*D*] crosses a critical value [*D*]_crit_. Below [*D*]_crit_, the attractor remains low-dimensional (few dominant 1-cycles), yielding low PE. Above [*D*]_crit_, the topology fragments—multiple loops emerge with dispersed lifetimes— raising PE. This curve provides *quantitative* evidence for a phase transition: a sharp PE increase indicates bifurcation of the global dynamical regime, operationally consistent with *ego dissolution* as Betti-1 stability breakdown. When experimental (true *𝜌*) and shuffled (*𝜌*_π_) conditions are compared, a larger PE increase in the experimental group supports receptor-specificity. Absence of a non-linear jump would falsify the bifurcation mechanism; flat or monotonic PE would indicate the system remains in a single dynamical regime across [*D*].

**Figure 3:**
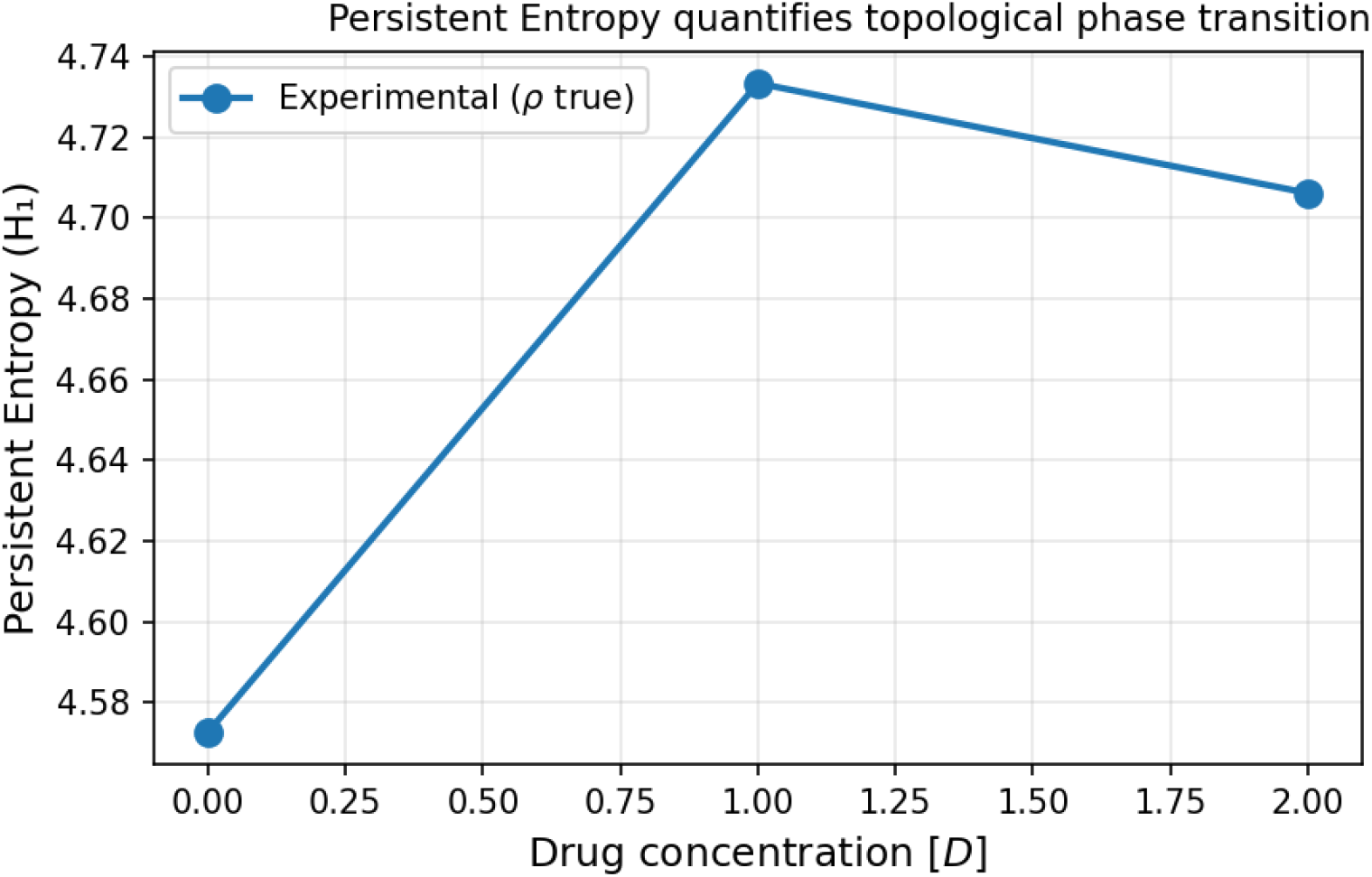
Persistent Entropy vs drug concentration, showing non-linear jump at critical value.

## Synthetic Parameter Table

| Parameter | Symbol | Value | Source |
| --- | --- | --- | --- |
| Parcellation | — | Schaefer 30 nodes | Schaefer et al., 2018 |
| Structural connectivity | $\mathbf{C}$ | HCP group-averaged | Van Essen et al., 2013 |
| Conduction velocity | $v$ | 5 m/s | Caminiti et al., 2009 |
| Baseline gain | $G_0$ | 1.0 | — |
| Receptor–gain coupling | $k$ | $[0.5, 5.0]$ | Bifurcation sweep |
| Noise amplitude | $\sigma$ | 0.02 | Euler–Maruyama |
| 5-HT2A receptor map | $\rho_i$ | Normalized PET | Beliveau et al., 2017; NeuroVault |
| Embedding dimension | $m$ | 3 | Takens |
| Embedding delay | $\tau$ | Data-dependent | Mutual information |
| E population time constant | $\tau_E$ | 10 ms | Wilson–Cowan |
| I population time constant | $\tau_I$ | 5 ms | Wilson–Cowan |
| E→E weight | $w_{EE}$ | 1.2 | — |
| I→E weight | $w_{IE}$ | 1.0 | — |
| E→I weight | $w_{EI}$ | 1.0 | — |
| I→I weight | $w_{II}$ | 0.7 | — |

## Discussion

### Topological Interpretation of State Transitions

The RCDT hypothesis interprets the shift from low-dimensional attractors to fragmented, high-dimensional structures as a **topological phase transition**. A single dominant Betti-1 cycle indicates constrained, quasi-periodic dynamics—consistent with a brain state that maintains coherent global structure. Fragmentation—multiple disconnected loops or higher homology—suggests decoupling of regional dynamics, reduced integrability, and potential correlates of *ego dissolution* when operationally defined as loss of low-dimensional Betti-1 stability.

This interpretation is mechanistic and geometric; it does not claim that topology *is* consciousness, but that topology may serve as an objective readout of dynamical regime changes induced by receptor-weighted pharmacology.

### Explicit Falsifiability

The RCDT hypothesis is rejected if any of the following obtain:

1. **Receptor-map independence**: Topological changes under drug perturbation are statistically indistinguishable when *𝜌*_*i*_ is replaced by a shuffled *𝜌*_𝜋(*i*)_. This would imply that receptor distribution is irrelevant—the effect would be generic gain modulation, not 5-HT2A-specific.
2. **Noise-equivalent topology**: The persistent homology of simulated time series is indistinguishable from that of appropriately scaled noise (e.g., surrogate data). This would invalidate the claim that deterministic structure is being detected.
3. **Monotonic or absent concentration–topology relationship**: Persistent entropy (or equivalent) shows no non-linear jump, or no systematic change, as [*D*] increases.

The hypothesis predicts a critical transition; its absence would falsify the bifurcation mechanism.

## Conclusion

We have formalized the Receptor-Constrained Dynamical Topology (RCDT) hypothesis—a mechanistic bridge from molecular receptor distribution to global topological manifold reorganization. The framework is biophysically grounded (Wilson–Cowan, structural connectivity, delays), pharmacologically explicit (5-HT2A-weighted gain), and topologically rigorous (Takens, Vietoris–Rips, persistent homology). Operational definitions (e.g., ego dissolution as Betti-1 breakdown) and explicit falsification criteria ensure that the hypothesis is testable. Future work will implement the proposed validation aims and compare predictions with empirical neuroimaging data.

## Supporting information

Supplementary material: Fig. S1 Receptor shuffling control; Fig. S2 Bifurcation sweep for parameter selection

## Data and Code Availability

- **5-HT2A receptor map**: NeuroVault (https://neurovault.org/)
- **Structural connectivity**: Human Connectome Project (https://www.humanconnectome.org/)
- **Simulation code**: https://github.com/lincNK/RCDT-Model

## References

Atienza, N. et al. (2016). Separating topological noise from features using persistent entropy. arXiv:1605.02885.

Beliveau, V. et al. (2017). A high-resolution in vivo atlas of the human brain’s serotonin system. Journal of Neuroscience, 37(1), 120–128.

Caminiti, R. et al. (2009). Evolution amplified processing with temporally dispersed slow neuronal connectivity in primates. Proceedings of the National Academy of Sciences, 106(46), 19551–19556.

Schaefer, A. et al. (2018). Local-global parcellation of the human cerebral cortex from intrinsic functional connectivity MRI. Cerebral Cortex, 28(9), 3095–3114.

Takens, F. (1981). Detecting strange attractors in turbulence. Lecture Notes in Mathematics, 898, 366–381.

Van Essen, D. C. et al. (2013). The WU-Minn Human Connectome Project. NeuroImage, 80, 62–79.

Wilson, H. R., & Cowan, J. D. (1972). Excitatory and inhibitory interactions in localized populations of model neurons. Biophysical Journal, 12(1), 1–24.

