## Supplementary figures and images for "Topological Phase Transitions in Whole-Brain Dynamics Driven by Spatially Heterogeneous Receptor Gain Modulation: A Receptor-Constrained Dynamical Topology (RCDT) Hypothesis"

### Supplementary material: Fig. S1 Receptor shuffling control; Fig. S2 Bifurcation sweep for parameter selection

#### Receptor Shuffling Control

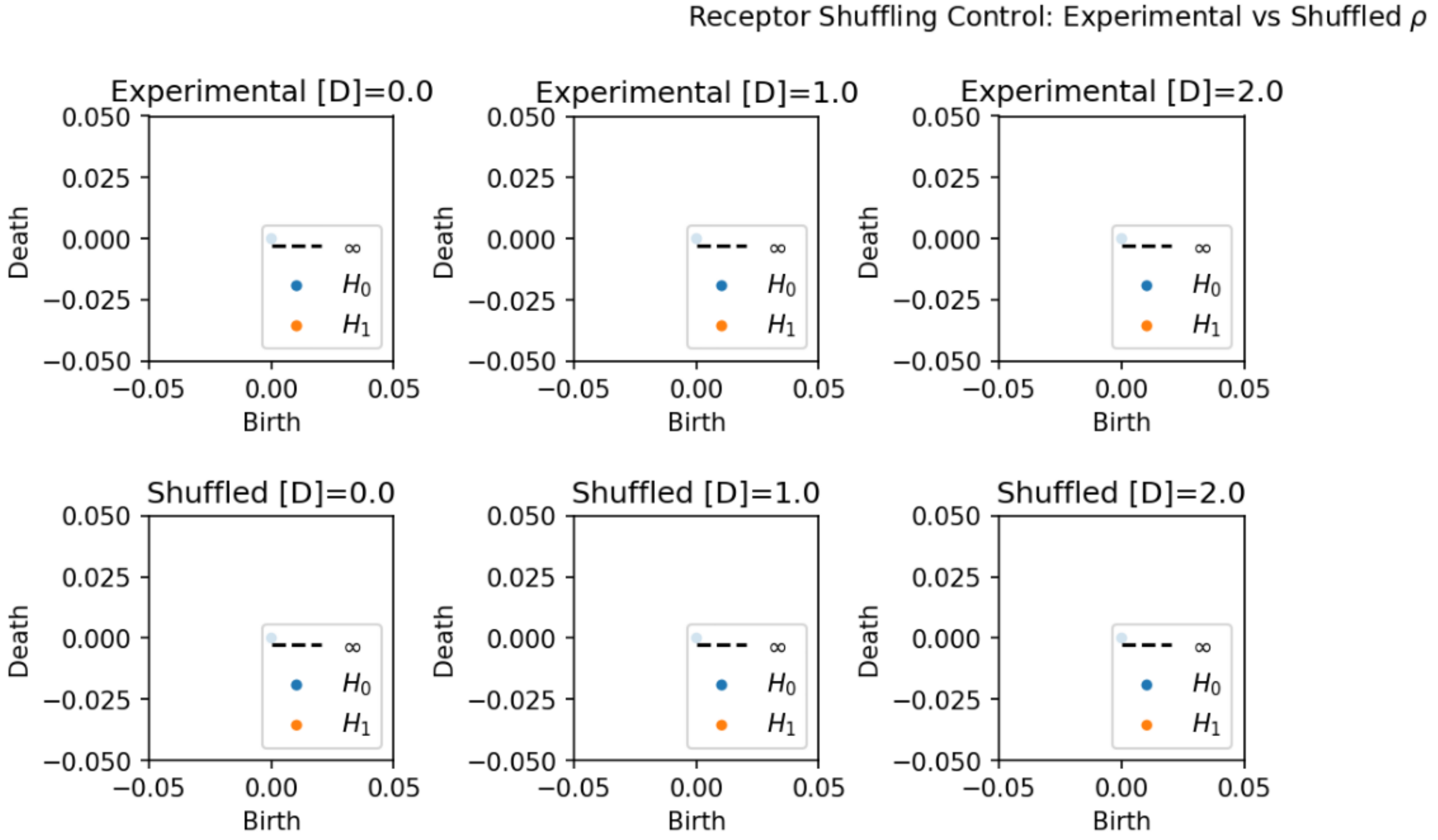


#### Bifurcation Sweep

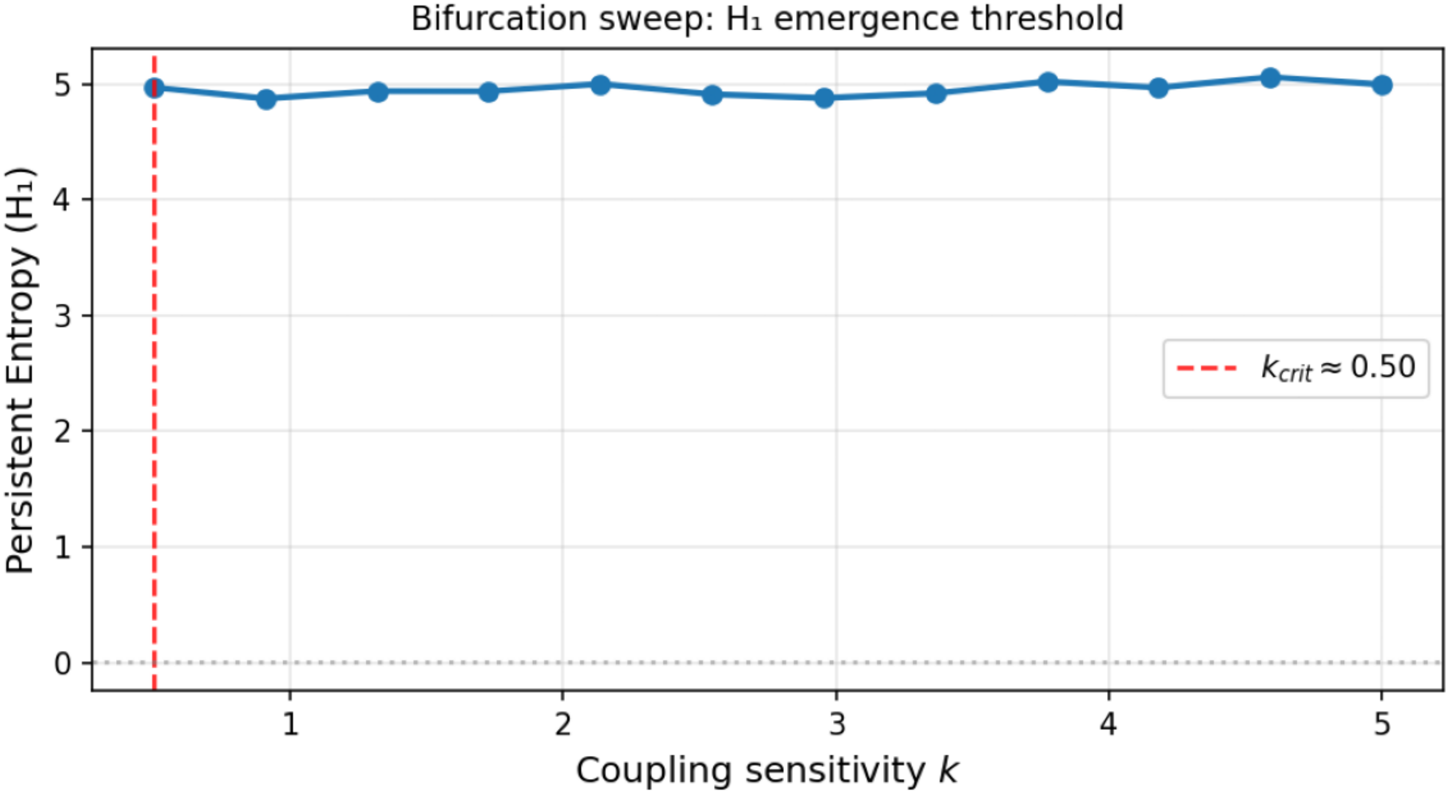
